# Cigarette smoke induces PPP1R15A via oxidative stress to modulate inflammatory cytokine production by bronchial epithelial cells

**DOI:** 10.1101/2024.12.24.630242

**Authors:** Gimano D. Amatngalim, Annemarie van Schadewijk, Anne M. van der Does, Maria C. Zarcone, Elke Malzer, Jayson Bradley, Jose A. Valer, Elaine Soon, Elizabeth Timmins, Mark Southwood, Maria G. Belvisi, Mark Birrell, Pieter S. Hiemstra, Stefan J. Marciniak

**Affiliations:** PulmoScience Lab, Department of Pulmonology, Leiden University Medical Center, Leiden, The Netherlands; Cambridge Institute for Medical Research (CIMR), University of Cambridge, Cambridge, UK; Division of Molecular Carcinogenesis, The Netherlands Cancer Institute, Amsterdam, the Netherlands; MRC Toxicology Unit, University of Cambridge, Cambridge, UK; Respiratory Pharmacology Group, National Heart & Lung Institute, Imperial College London; Royal Papworth Hospital NHS Foundation Trust, Cambridge, UK

**Author notes:** Joint senior authors (Pieter S. Hiemstra, Stefan J. Marciniak).

**Keywords:** smoke, PPP1R15A, lung, airway epithelium, oxidative stress

## Abstract

Cigarette smoke is the principal cause of chronic obstructive pulmonary disease (COPD) in Western societies. It induces pulmonary inflammation and activates multiple stress signalling pathways in bronchial epithelial cells, including the integrated stress response (ISR). The ISR is triggered by phosphorylation of the translation initiation factor eIF2α in response to a variety of insults and is functionally antagonised by the eIF2α phosphatase subunit PPP1R15A. Deletion of PPP1R15A exaggerates the ISR and, in many models, protects against cellular stress. However, loss of PPP1R15A was also shown to exaggerate pulmonary fibrosis in bleomycin-challenged mice. We therefore wished to understand the role of PPP1R15A in the pulmonary response to cigarette smoke, which can cause both COPD and pulmonary fibrosis. Whole cigarette smoke rapidly activated the ISR in differentiated mucociliary human primary bronchial epithelial cells, with those from COPD patients exhibiting higher levels of ISR. Acute exposure to cigarette smoke condensate induced a pro-inflammatory cytokine response accompanied by PPP1R15A expression. In this model, PPP1R15A induction was independent of eIF2α phosphorylation but was suppressed by antioxidants, suggesting a response to oxidative stress. In vivo studies in mice revealed that PPP1R15A deficiency enhanced the secretion of proinflammatory cytokines in mice, suggesting a regulatory role for PPP1R15A in modulating the inflammatory response to smoke. Taken together, these data suggest that PPP1R15A is induced by cigarette smoke by an ISR-independent oxidative stress mechanism and ameliorates the pro-inflammatory cytokine response to cigarette smoke.

## Introduction

The airway epithelium defends against respiratory viruses and bacteria through its barrier function, mucociliary clearance, and innate immune defence mechanisms (1–4). Environmental stressors, including pathogens and toxic fumes, can damage the airway epithelium to impair these protective processes (5). Mechanisms have therefore evolved to adapt airway epithelial cells to stress, for example through detoxification by metabolic enzymes and the generation of antioxidants (6).

In Western societies, the primary cause of chronic obstructive pulmonary disease (COPD) is cigarette smoking (7). Exposure to cigarette smoke triggers several stress-responsive pathways, including the unfolded protein response (UPR) to endoplasmic reticulum (ER) stress (8–11). Cigarette smoke extract preferentially activates the PERK-mediated arm of the UPR in the bronchial epithelial cell line BEAS-2B through oxidative stress (12). PERK is an ER stress-sensing kinase that phosphorylates the α subunit of eukaryotic translation initiation factor 2 (eIF2α) (13). Since eIF2α can also be phosphorylated by kinases that respond to a variety of different stresses, including GCN2 responding to ribosome stalling, HRI to iron deficiency and PKR to viral double-stranded RNA, the pathway downstream has been named the integrated stress response (ISR) (14). Phosphorylation of eIF2α is cytoprotective in part through limiting new protein synthesis and through induction of ISR target genes downstream of the transcription factors ATF4, ATF3 and CHOP (15, 16). Interestingly, the gene expression signature present in bronchial tissue from COPD patients is enriched for genes regulated by ATF4 (15). Together, ATF4 and CHOP eventually induce PPP1R15A (also known as GADD34), which dephosphorylates eIF2α to terminate the ISR (17). However, during sustained stress, the recovery of protein translation mediated by PPP1R15A can be toxic (17). PPP1R15A is therefore a key regulator of cell viability in cells exposed to stress.

*In vitro* studies with cell lines have shown UPR and ISR activation by cigarette smoke, but relatively little is known about activation of the UPR and ISR in differentiated primary airway epithelial cells or tissues. Moreover, the mechanisms underlying cigarette smoke-mediated activation of the ISR in such lung cells are incompletely understood.

Previously, we showed that brief exposure of airway epithelial cells to whole cigarette smoke had a mild cytotoxic effect, while cells displayed transient inflammatory responses and a temporary impairment in the epithelial barrier integrity (18). Furthermore, more recently we demonstrated that the transcriptional responses of the airway epithelial cells in this whole cigarette smoke exposure model mirror those of in vivo exposure of the human airway epithelium (19). Therefore, in the present study, we examined activation of the ISR in airway epithelial cells *in vitro* and *in vivo* by cigarette smoke, and we also compared the responses of human cell cultures derived from individuals with COPD and non-COPD controls.

We show that cigarette smoke increases the expression of PPP1R15A via a non-ISR oxidative stress-dependent pathway. In mice exposed to cigarette smoke for two weeks, genetic absence of PPP1R15A led to increased production of proinflammatory cytokines IL-1, KC and IL-17. These findings suggest that PPP1R15A is induced by multiple pathways to modify the airway inflammatory response to smoke.

## Materials and methods

### Cell culture

Primary bronchial epithelial cells (PBEC) were isolated from tumour-free resected lung tissue obtained during surgery for lung cancer at the Leiden University Medical Center, and cultured at the air-liquid interface (ALI) or submerged conditions (S). Initially, lung tissue samples were enrolled in the biobank via a no-objection system for coded anonymous further use of such tissue (www.coreon.org). Patients from whom lung tissue was derived were enrolled in the biobank via a no-objection system for coded anonymous further use of such tissue. Samples from this Biobank were approved for research use by the institutional medical ethical committee (BB22.006/AB/ab). Culturing of ALI-PBEC and S-PBEC was performed as previously reported (18). In ALI-PBEC cultures, cells were seeded on 0.4 µm pore sized semi-permeable transwell membranes (Corning Costar, Cambridge, MA, USA) and first cultured in submerged conditions in a 1:1 mixture of bronchial epithelial growth medium (BEGM) (Lonza, Verviers, Belgium) and Dulbecco’s modified Eagle’s medium (DMEM) (Gibco, Bleiswijk, The Netherlands) containing 1 mM HEPES (Lonza), 100 U/mL penicillin and 100 µg/ml streptomycin (Lonza) (hereafter referred to as B/D culture medium), supplemented with SingleQuot BEGM supplements (except Gentamycin) (Lonza), 15 ng/ml retinoic acid (Sigma-Aldrich) and 1 mg/ml BSA (Sigma-Aldrich). When monolayers were confluent, the apical medium was removed, and cells were cultured in air-exposed conditions for at least 2 weeks to allow mucociliary differentiation. In western blot experiments, cells were cultured overnight and during the experiment with starvation medium, consisting of B/D medium without BSA and the SingleQuot supplements BPE and EGF. ALI-PBEC cultures were used from COPD patients and non-COPD (ex)smokers, for whom the disease status was determined based on pre-surgery lung function data according to the Global Initiative for Chronic Obstructive Lung Disease classification (7) (Table 1). S-PBEC were cultured in regular tissue culture plates with B/D culture medium as earlier described, but without the additional 15 ng/ml retinoic acid supplementation. Cells were used when approximately 80 -90% confluent. In all experiments, S-PBEC were cultured overnight prior to the experiment and during the experiment with starvation medium, which was similar to ALI-PBEC starvation medium, but lacked the additional 15 ng/ml retinoic acid supplementation. In addition to the PBEC cultures, also eIF2α^AA^ and eIF2α^SS^ mouse embryonic fibroblasts (MEFs) were used, and cultured as previously described (20).

**Table 1:**
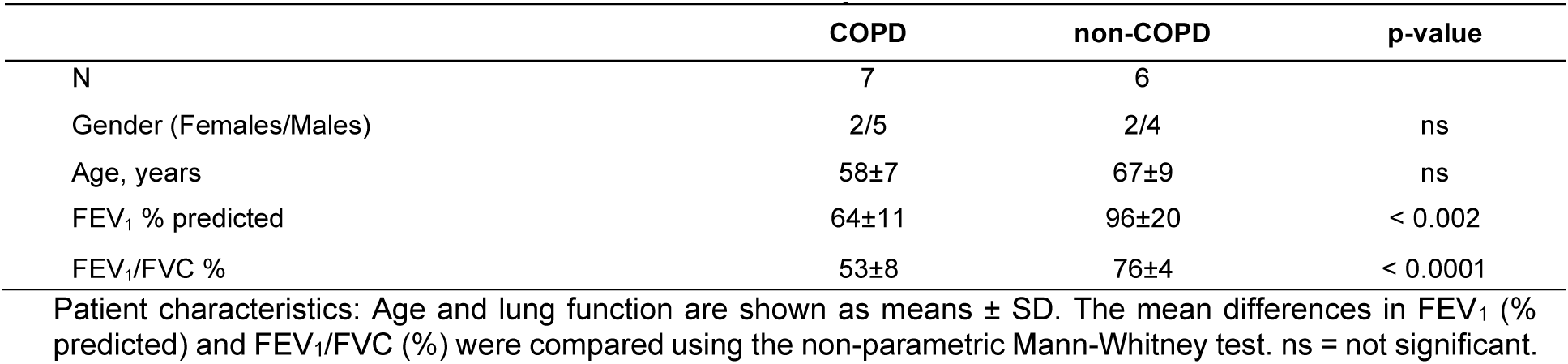
Characteristics of COPD and non-COPD patients.

|  | **COPD** | **non-COPD** | **p-value** |
|---|---|---|---|
| N | 7 | 6 |  |
| Gender (Females/Males) | 2/5 | 2/4 | ns |
| Age, years | 58±7 | 67±9 | ns |
| FEV<sub>1</sub> % predicted | 64±11 | 96±20 | < 0.002 |
| FEV<sub>1</sub>/FVC % | 53±8 | 76±4 | < 0.0001 |

Patient characteristics: Age and lung function are shown as means ± SD. The mean differences in FEV<sub>1</sub> (% predicted) and FEV<sub>1</sub>/FVC (%) were compared using the non-parametric Mann-Whitney test. ns = not significant.

### Cigarette smoke (CS) exposure bronchial epithelial cell model and other stimuli

ALI-PBEC cultures were exposed to whole cigarette smoke (CS) using an exposure model previously described (18). In short, cells were exposed to whole smoke derived from one standard research cigarette (3R4F), after which cells were incubated for the indicated periods of time. S-PBEC and MEF cultures were treated with cigarette smoke condensate (CSC), which was prepared as previously reported (21). Cells were treated with CSC via a “pulse” method described in Figure 4 and corresponding text in the Results section, similar to the whole CS exposure protocol. In indicated experiments, cells were pre-treated for 1 hour and treated during the experiment with the antioxidant N-acetylcysteine (NAC) (Sigma). As indicated, in some experiments cells were treated with thapsigargin (Tg) or tunicamycin (Tm) (both Sigma) as positive controls for ER stress.

### Whole body cigarette smoke (CS) mouse model

PPP1R15A-deficient (*Ppp1r15a^ΔC/ΔC^*) mice, generated (22) and backcrossed to 97% purity C57BL/6 as described, and wildtype controls were exposed to whole cigarette smoke using a model previously reported and characterised (23). Briefly, CS was generated from 3R4F cigarettes (filters removed; Tobacco Health Research Institute, University of Kentucky, Lexington, KY, USA) and drawn through an exposure chamber (136 L capacity; Teague Enterprises, CA, USA) in which a fan was placed to ensure even distribution. Exposures at 500 ml/min lasted 50 minutes, followed by a 10-minute venting period to flush the chamber. Total smoke particulate (TSP) was assessed every 15 minutes using a TSP sampling unit (Teague Enterprises). Mice were exposed twice daily for 3 or 14 days. Lung tissue for RT-PCR analysis of mRNA levels and bronchoalveolar lavage fluid (BALF) for cytokine measurements were collected 24 hours after the last exposure and processed as described. Animal studies were conducted in accordance with the UK Animals (Scientific Procedures) Act 1986 under the Home Office project licenses 5789170 and 70/7212. Animals were housed with free access to food and water, and all procedures complied with institutional and national guidelines for animal welfare.

### Western blot

Cell lysates were prepared in Harvest Buffer (Buffer H), consisting of 10 mM HEPES pH 7.9, 50 mM NaCl, 0.5 mM sucrose, 0.1 mM EDTA, 0.5% (v/v) Triton X-100, 1 mM DTT, Protease inhibitor cocktail (Roche Applied Science, Mannheim, Germany), phosphatase inhibitors (10 mM tetrasodium pyrophosphate, 17.5 mM β-glycerophosphate, and 100 mM NaF), and 1 mM PMSF. Nuclear extraction was performed using the NE-PER Nuclear Protein Extraction Kit (Thermo Scientific, Rockford, IL) according to the manufacturer’s protocol. Cell lysates were diluted in 4x sample buffer consisting of 0.2 M Tris-HCl pH 6.8, 16% [v/v] glycerol, 4% [w/v] SDS, 4% [v/v] 2-mercaptoethanol and 0.003% [w/v] bromophenol blue. Samples were separated using a 10% SDS-PAGE gel, and proteins were transferred onto nitrocellulose membrane. Primary antibodies were used against: eIF2α, phospho-eIF2α, ATF4, TBP (Cell Signaling Technology), PPP1R15A/GADD34 (ProteinTech) and puromycin (Millipore) (all 1:1000 diluted). After secondary antibody incubation, protein detection was visualised using ECL (ThermoScientific) or the LI-COR Odyssey Infrared Imaging System (LI-COR Biosciences). Quantification of protein bands was done by densitometry using ImageJ software (National Institutes of Health, Bethesda, MD, USA).

### qPCR

*Cell culture CS model.* RNA was isolated with the Maxwell tissue RNA extraction kit (Promega) or Qiagen RNeasy mini kit (Qiagen). cDNA synthesis and qPCR were performed as previously described (see Table 2 for qPCR primer pairs) (18, 20). Relative gene expression was calculated according to the standard curve method. ATP5B and RPL13A were selected using the “Genorm method” (Genorm; Primer Design, Southampton, UK) as reference genes for PBEC. β-actin was used as a reference gene for MEFs. *In vivo CS model*. RNA was isolated from flash-frozen lung tissue using the Qiagen RNeasy Universal Kit (Qiagen), then analysed with the same protocol as used for cell lines.

**Table 2:**
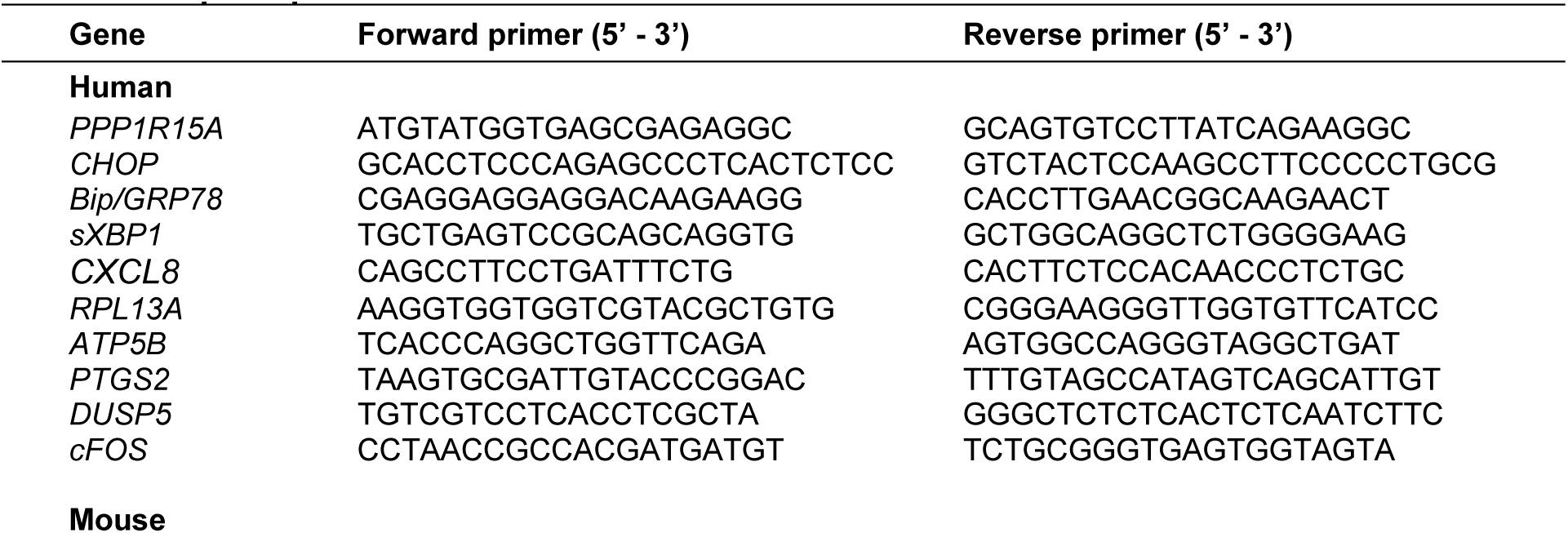
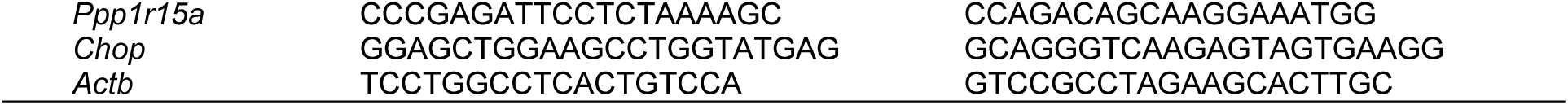
qPCR primers.

### ELISA & LDH release assay

Protein secretion of IL-8/CXCL8 (R&D Systems) and LDH release (Roche) were determined in cell culture supernatants according to the manufacturers’ instructions. BALF cytokines from smoke-exposed mice were measured using a Mesoscale Discovery (MSD) multiplex assay. BALF samples (500 μL) were concentrated approximately five-fold by centrifugation through 3 kDa molecular weight cut-off Amicon filters to a final volume of approximately 100 μL before analysis. The assay measured TNF-α, GM-CSF, IL-1β, IL-6, IL-10, IL-17, KC, MCP-1, MIP-1α and MIP-2. Cytokine concentrations were analysed without correction for the concentration factor.

#### Analysis of published mouse lung single-cell RNA sequencing data

Our single-cell RNA sequencing of murine lung tissues was published previously and made openly accessible at the Gene Expression Omnibus database with the reference <u>GSE294702</u> (24). This was re-analysed focusing on *Ppp1r15a* and *Ppp1r15b*.

### Immunohistochemistry for PPP1R15A

Lungs from 3-month-old littermate wild-type and *Ppp1r15a^ΔC/ΔC^*mice were inflation-fixed, embedded in paraffin, sectioned, and subjected to immunohistochemistry. Sections were incubated with rabbit anti-PPP1R15A/GADD34 antibody (ProteinTech; 1:200 dilution) followed by detection using a standard horseradish peroxidase-based immunohistochemical method. Immunoreactivity was visualised with DAB chromogen, and sections were counterstained with haematoxylin.

### Statistics

Results were analysed using GraphPad Prism 6.0 (GraphPad Inc). As indicated in the figures, statistical tests used for data analysis were (un)paired t-test or 2-way ANOVA, with post-hoc Bonferroni correction for multiple analyses. Differences with a p-value < 0.05 were considered statistically significant.

## Results

### Whole cigarette smoke exposure increases the ISR in ALI-PBEC

We first examined the effect of whole cigarette smoke (CS) on activation of the integrated stress response (ISR) in mucociliary differentiated human primary bronchial epithelial cells (ALI-PBEC). Cell cultures were exposed to whole smoke from a single cigarette (Figure 1A), which in previous studies was shown to cause an acute and transient disruption of the epithelial barrier, impairment of wound repair and induction of innate immune responses with minimal cytotoxic effects (18, 25, 26). Acute activation of the ISR was detected with phosphorylation of eIF2α and nuclear accumulation of ATF4 within the first 2 hours after exposure (Figure 1B, C). Time-dependent induction of expression of the ISR genes *PPP1R15A* and *CHOP* was observed in CS-exposed cells, progressively up to 2 hours following exposure (Figure 1D). As expected for an activator of the ISR, cigarette smoke caused transient phosphorylation of eIF2α with an associated inhibition of protein synthesis, which both abated as PPP1R15A protein accumulated (Figure 2A-C).

**Figure 1.**
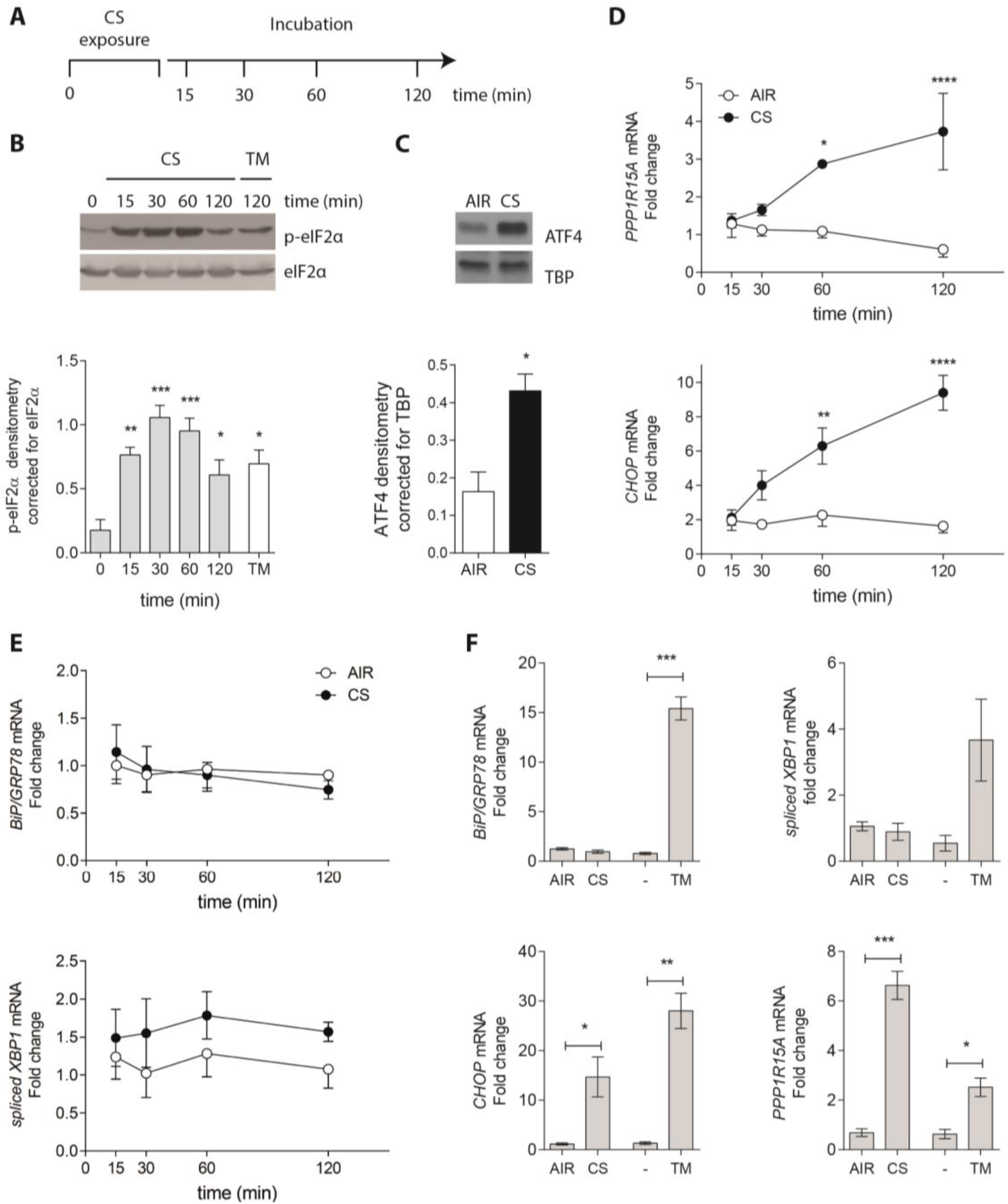
Effect of whole cigarette smoke (CS) exposure on ISR-activation in ALI-PBEC cultures. **A)** Schematic of the whole CS exposure experiment, in which ALI-PBEC were exposed to CS from a single cigarette, followed by incubation for different time periods ranging from 15 - 120 min. **B)** eIF2α phosphorylation was determined following 15-120 min of whole CS exposure. The ER stress inducing agent tunicamycin (TM) was used as positive control. Results were quantified by densitometry using total eIF2α as loading control. **C)** Nuclear localization of ATF4 was determined 2 h after CS exposure. Results were quantified by densitometry using TATA binding protein (TBP) as loading control. **D)** Analysis of *PPP1R15A* and *CHOP,* **E)** *BiP* and *sXBP1* mRNA expression 15-120 min after CS exposure. **F)** mRNA expression of *PPP1R15A*, *CHOP, BiP* and *sXBP1* after Air/CS exposure or TM after 6 h incubation period. Results are shown as mean ± SEM of at least 3 different donors. Analysis of differences was conducted using (B,D,E) a one-way ANOVA with a Bonferroni *post-hoc* test and (C,F) a paired t-test. * p < 0.05, ** p < 0.01, *** p < 0.001.

**Figure 2.**
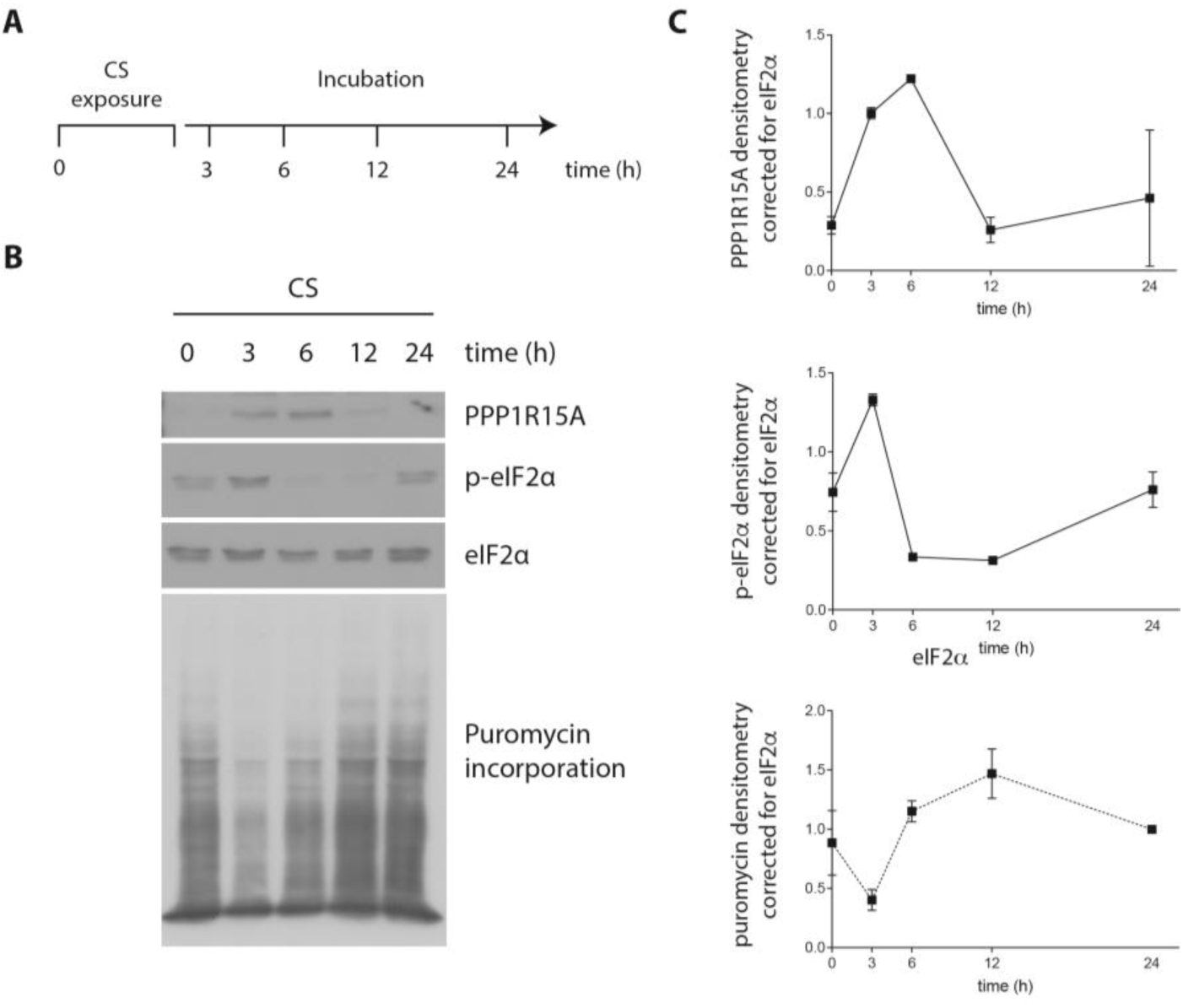
Effect of whole CS exposure on protein translation in ALI-PBEC cultures. **A)** Schematic of the experiment, in which ALI-PBEC were exposed to CS from a single cigarette, followed by incubation for different time periods ranging from 0-24 h. **B)** Analysis of PPP1R15A protein expression, eIF2α phosphorylation, and puromycin incorporation as marker of protein translation. **C)** Results were quantified by densitometry using total eIF2α as loading control. Experiments were conducted in 2 independent donors showing similar results.

Previous studies suggested the activation of the ISR by cigarette smoke was a response to ER stress (12). However, in stark contrast to the smoke-induced increase in *CHOP* and *PPP1R15A*, we did not observe increased expression of the ER chaperone *BiP* nor splicing of *XBP1* mRNA, a proximal mediator of the UPR downstream of the ER stress sensor IRE1 (Figure 1E&F). Tunicamycin served as a positive control for ER stress and robustly induced *BiP* and *XBP1* mRNA splicing in ALI-PBEC cultures (Figure 1E&F).

### COPD airway epithelial cells display higher CS-induced expression of CHOP and PPP1R15A

Next, we examined the induction of the *CHOP* and *PPP1R15A* mRNA in ALI-PBECs from COPD patients and (ex)smokers who had not developed COPD. In line with our earlier observations, cigarette smoke induced the expression of *CHOP* and *PPP1R15A* soon after exposure, returning to baseline over 24 hours (Figure 3A). Remarkably, the early induction of *PPP1R15A* and *CHOP* was significantly higher in COPD airway epithelial cultures compared with non-COPD controls. Consequently, induction of PPP1R15A and CHOP at 3 h showed a significant negative correlation with both FEV_1_ (Figure 3B) and FEV_1_/FVC (Figure 3C). By contrast, other genes thought to be upregulated in COPD lungs by ATF4, including *PTGS2*, *DUSP5* and *cFOS* (15), were transiently induced by cigarette smoke but did not differ between COPD and non-COPD ALI-PBECs (Supplemental Figure 1).

**Figure 3.**
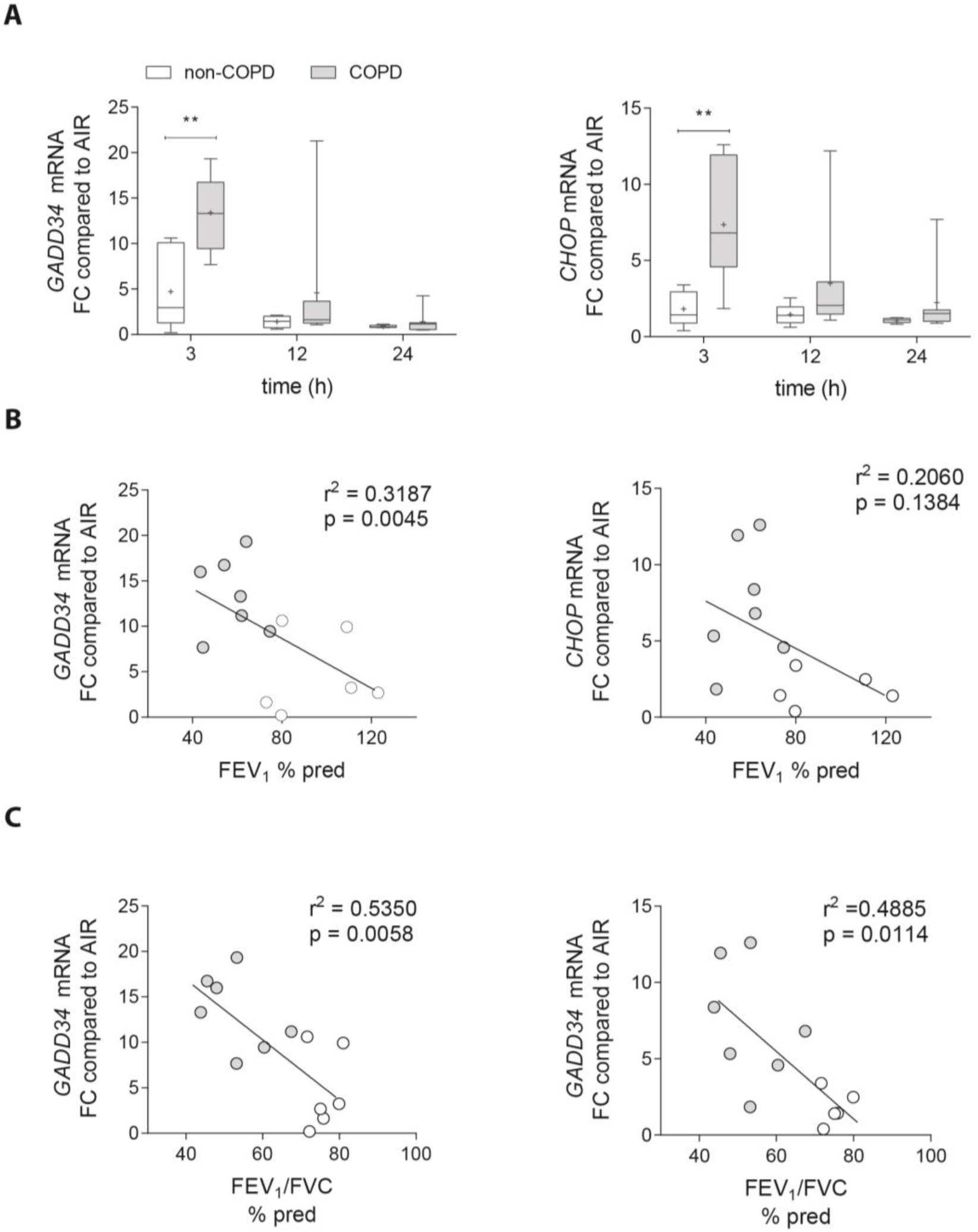
**CS-induced expression of *PPP1R15A* and *CHOP* in COPD and non-COPD ALI-PBEC**. **A)** ALI-PBEC from COPD and non-COPD donors were exposed to whole CS and incubated for 3,12, and 24 h after which *PPP1R15A* and *CHOP* mRNA expression was determined and expressed as fold-change (FC) compared to air-exposed cultures. **B)** Correlation between fold change in mRNA expression compared to air controls of *PPP1R15A* and *CHOP* mRNA, with FEV_1_ (% pred), and **C)** FEV_1_/FVC (% pred). n=7 COPD and n=6 non-COPD donors were used in the analysis. Analysis of differences was conducted using a one-way ANOVA with a Bonferroni *post-hoc* test. ** p < 0.01. Correlations were assessed by determining the linear regression.

**Figure 4.**
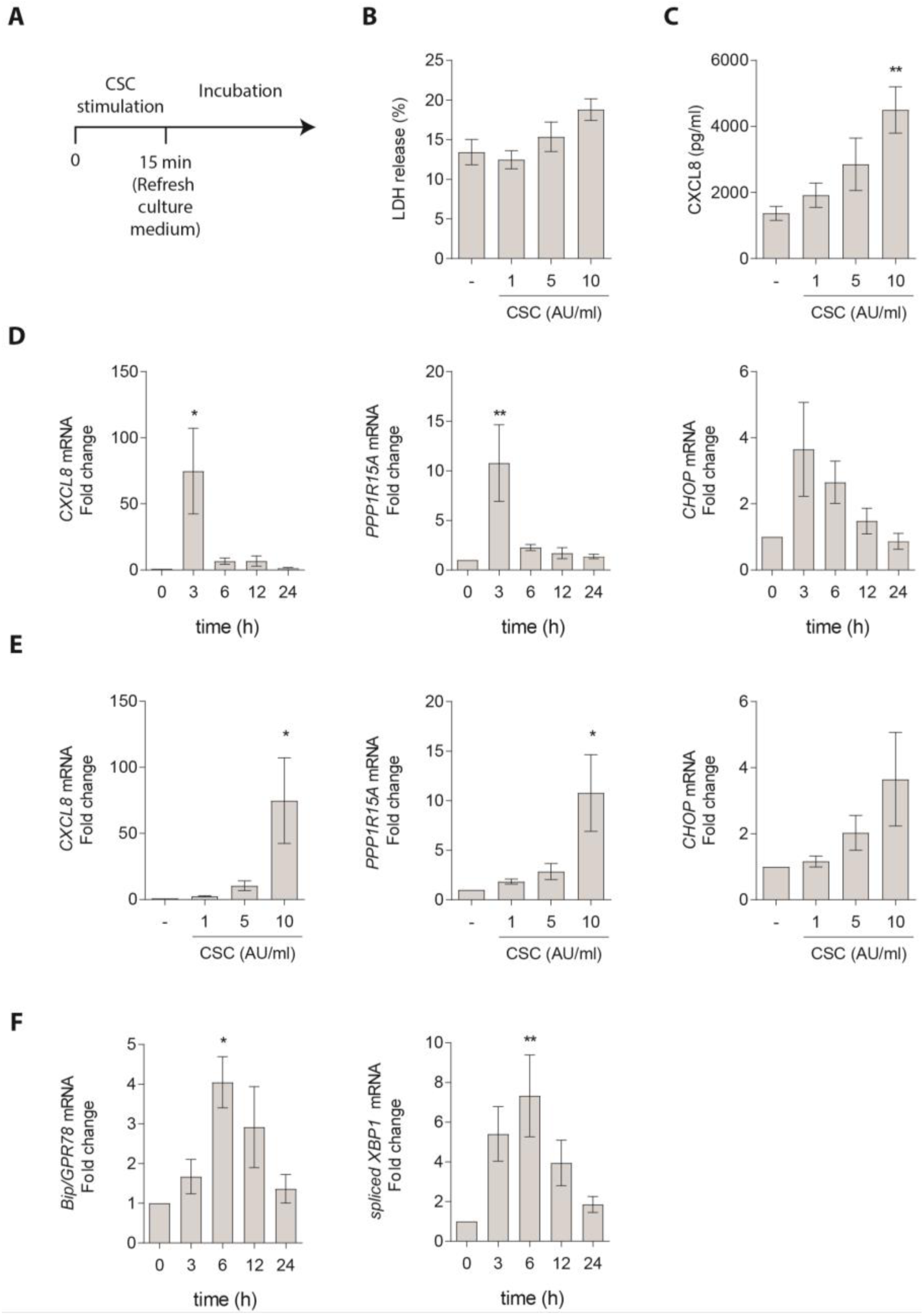
Effect of CSC on ISR activation in undifferentiated S-PBEC. **A)** Schematic of experiments with CSC pulse exposure of S-PBEC cultures. **B)** Assessment of the cytotoxic effect of 1-10 AU/ml CSC on S-PBEC after 24 h incubation. Percentage LDH release was used as measurement of cytotoxicity. **C)** Assessment of CXCL8 protein release in the cell culture supernatant of S-PBEC stimulated with 1, 5 and, 10 AU/ml CSC and incubated for 24 h. **D)** Assessment of *CXCL8*, *PPP1R15A* and *CHOP* mRNA expression at different time points (3-24 h) after 10 AU/ml CSC exposure of S-PBEC. mRNA expression is shown as fold change compared to control-treated cells. **E)** Assessment of *CXCL8*, *PPP1R15A* and *CHOP* mRNA expression in S-PBEC after stimulation with 1, 5, and 10 AU/ml CSC exposure and 3 h incubation. mRNA expression is shown as fold change compared to control-treated cells. **F)** mRNA expression of *BiP* and *sXBP1* at different time points (3-24 h) after 10 AU/ml CSC exposure of S-PBEC. mRNA expression is shown as fold change compared to controls. Results are shown as mean ± SEM using cells of at least 3 independent donors. Analysis of differences was conducted using a one-way ANOVA with a Bonferroni *post-hoc* test. * p < 0.05, ** p < 0.01..

### Cigarette smoke extract increases the ISR in submerged cultured PBEC

After showing that exposure to whole cigarette smoke activates the ISR in well-differentiated ALI-PBEC cultures, we next employed simpler, conventional submerged cell cultures of PBEC (S-PBEC). In view of the submerged nature of the culture, we examined the effect of acute exposure to cigarette smoke condensate for 15 minutes instead of whole cigarette smoke (Figure 4A). The reasons for this approach were (i) to verify our findings in a different culture model with a different type of smoke exposure, and (ii) as preparation for the mechanism-focused experiments with fibroblasts described in the next paragraph. In contrast to the marked toxicity of prolonged exposure (21), pulsed exposure caused only a non-significant small increase in LDH release (Figure 4B). However, even brief exposure to cigarette smoke condensate caused a dose-dependent increase in protein secretion of the neutrophil chemoattractant IL-8/CXCL8 (Figure 4C), which was accompanied by a rapid and dose-dependent induction of *CXCL8* mRNA (Figure 4D, E). Consistent with our previous studies with whole cigarette smoke (18), pulsed cigarette smoke condensate is pro-inflammatory with minimal immediate cytotoxicity.

Similar to *CXCL8*, induction of *PPP1R15A* and *CHOP* mRNA expression was also rapidly and transiently induced in a dose-dependent manner by cigarette smoke condensate, peaking 3 hours following exposure (Figure 4D, E). In contrast to our observations with whole cigarette smoke-exposed differentiated ALI-PBECs, cigarette smoke condensate also induced mRNA expression of *BiP* and spliced *XBP1*, peaking at 6 h after exposure (Figure 4F). This might plausibly reflect differences in the constituents or concentration of condensate versus whole smoke (27, 28), or differences in the response of cultures consisting only of basal epithelial cells (S-PBEC), versus mucociliary differentiated ALI-PBEC cultures.

### Cigarette smoke condensate induces PPP1R15A independently of eIF2α phosphorylation

To gain mechanistic insight into the induction of *PPP1R15A* by cigarette smoke condensate, we used wildtype (eIF2α^SS^) mouse embryonic fibroblasts (MEFs) and MEFs incapable of eIF2α phosphorylation owing to the biallelic mutation of eIF2α serine 51 (eIF2α^AA^) (29). As was seen with PBECs, cigarette smoke condensate induced expression of both *Chop* and *Ppp1r15a* in eIF2α^SS^ MEFs (Figure 5A). Unexpectedly, *Ppp1r15a* induction was preserved in MEFs in which the ISR had been disabled (eIF2α^AA^), whereas *Chop* induction was markedly attenuated. Indeed, in eIF2α^AA^ MEFs, cigarette smoke condensate induced PPP1R15A at a concentration lower than that required in wild-type cells, suggesting an increased sensitivity to this stress. Although the eIF2α^AA^ mutation completely abrogated the phosphorylation of eIF2α in response to cigarette smoke condensate, the protein expression of PPP1R15A was similar in both eIF2α^AA^ and eIF2α^SS^ MEFs following cigarette smoke condensate (Figure 5B).

**Figure 5.**
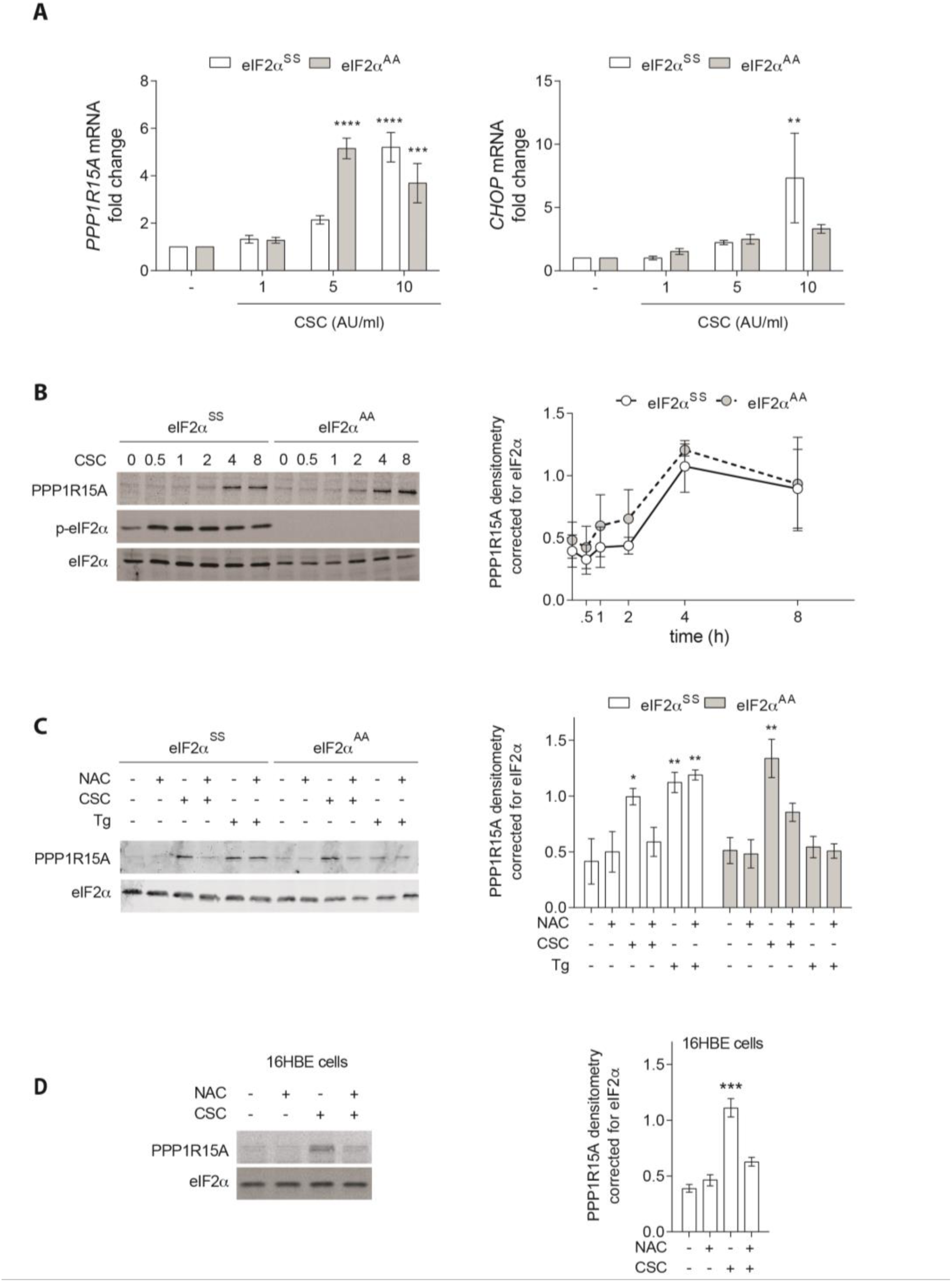
**ISR-independent expression of PPP1R15A in cigarette smoke condensate (CSC) exposure**. **A)** Assessment of *PPP1R15A* and *CHOP* mRNA expression in eIF2α^AA^ and eIF2α^SS^ MEFs stimulated with 1.5 or 10 AU/ml CSC after pulse stimulation and followed by 24 h incubation. Results are shown as fold change in mRNA compared to control-treated cells. **B)** Assessment of PPP1R15A protein expression in eIF2α^AA^ and eIF2α^SS^ MEFs stimulated with 10 AU/ml CSC after pulse stimulation and followed by incubation at different time points. Results were quantified by densitometry using total eIF2α as loading control. **C)** Assessment of the effect of N-acetyl cysteine (NAC) on CSC and thapsigargin (Tg)-induced expression of PPP1R15A in eIF2α^AA^ and eIF2α^SS^ MEFs after 8 h incubation. **D)** Immunoblot of PPP1R15A protein expression in 16HBE cells treated with 10 AU/ml CSC for 8 h ± N-acetyl cysteine (NAC). Results were quantified by densitometry using total eIF2α as loading control. Results are shown as mean ± SEM of at least 3 repeats. Analysis of differences from control was conducted using a one-way ANOVA with a Bonferroni *post-hoc* test. * p < 0.05, ** p < 0.01, *** p < 0.001.

It is known that ISR-deficient cells are impaired in their response to oxidative stress (9, 30–33). It is also known that altered redox plays an important role in the cytotoxic effects of cigarette smoke (34–37). We therefore wondered whether the ISR-independent expression of PPP1R15A might involve oxidative stress. To test this, eIF2α^AA^ and eIF2α^SS^ MEFs were treated with the antioxidant N-acetylcysteine (NAC) and cigarette smoke condensate individually and in combination. Cigarette smoke condensate-induced expression of PPP1R15A was inhibited by NAC in both eIF2α^AA^ and eIF2α^SS^ MEFs (Figure 5C). Similarly, in the airway epithelial cell line 16HBE, CSC induced PPP1R15A expression, and this was abrogated by cotreatment with NAC (Figure 5D).

Thapsigargin, which served as a control for ER stress-induced activation of the ISR, also induced expression of PPP1R15A, but only in ISR-competent eIF2α^SS^ MEFs, and this induction was insensitive to NAC (Figure 5C). These findings indicate that cigarette smoke condensate can induce PPP1R15A via an ISR-independent oxidative stress pathway.

### PPP1R15A is expressed in mouse lung

PPP1R15A is regarded as a stress-induced eIF2α phosphatase, with PPP1R15B serving as the constitutively expressed paralog (17, 38–40). Since most previous studies of PPP1R15A had focused on cultured cell lines, we examined its expression in lung tissue from mice without physiological stress. Using our published single-cell transcriptomic data from mouse lung (24), we found PPP1R15A, rather than PPP1R15B, to be the most highly expressed of the two phosphatase subunits in several lung cell types, including ciliated epithelial cells (Figure 6A). We validate this at the protein level by immunohistochemistry, with PPP1R15A-deficient (*Ppp1r15a^ΔC/ΔC^*) mice serving as a negative control (Figure 6B). Anti-PPP1R15A staining was detectable in airway and alveolar epithelial cells, while the corresponding staining was markedly reduced in *Ppp1r15a^ΔC/ΔC^* mice, which express an unstable C-terminal truncation (17, 41)

**Figure 6.**
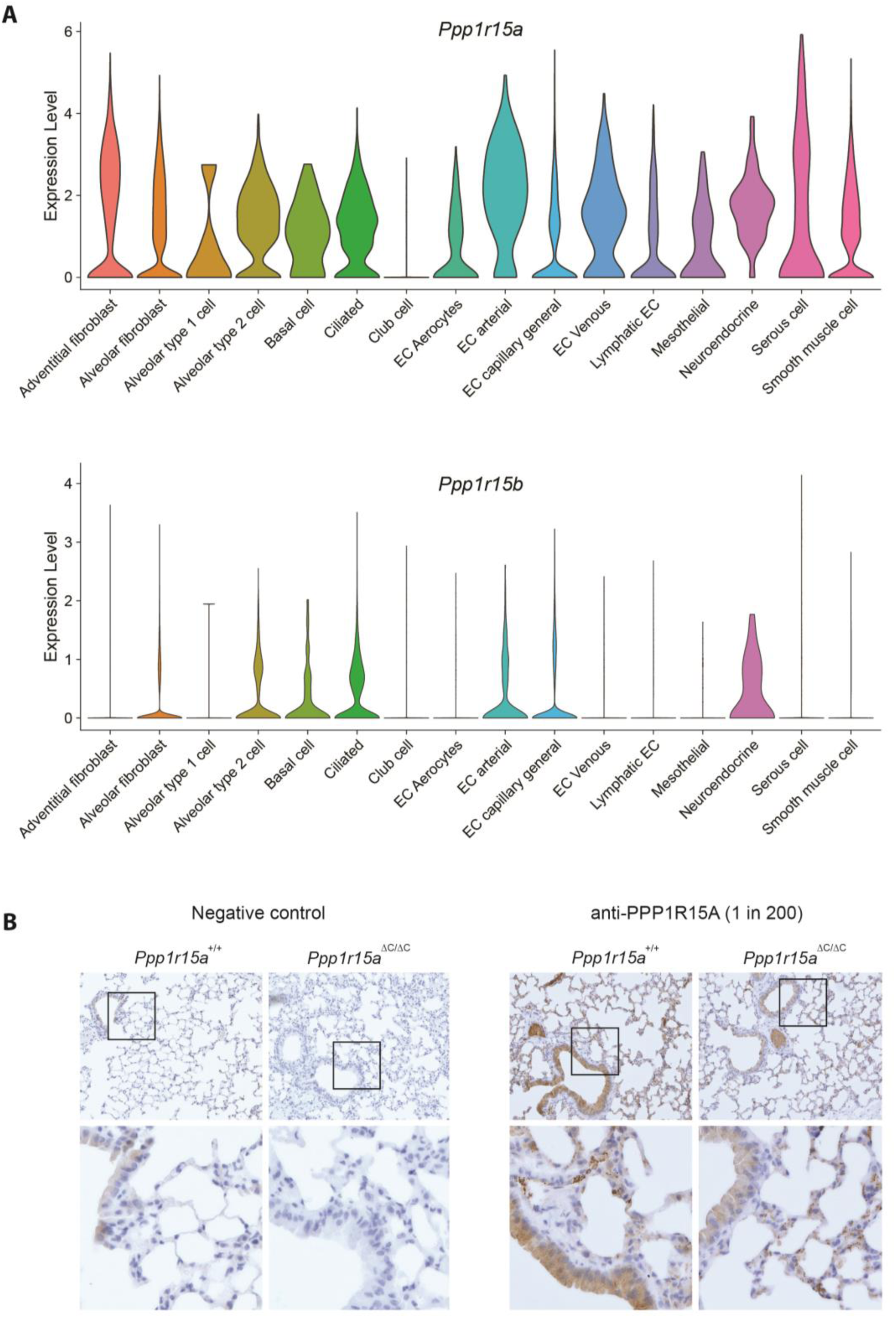
PPP1R15A is expressed in the mouse lung. (A) Violin plots of *Ppp1r15a* and *Ppp1r15b* RNA expression in wild-type mouse lung tissue, using published data (24). (B) Representative immunohistochemical staining for PPP1R15A in lungs from 3-month-old wild-type and *Ppp1r15a^ΔC/ΔC^* littermate mice. Brown staining indicates PPP1R15A-positive cells. Staining is evident in airway and alveolar epithelial cells of wild-type mice and is markedly reduced in *Ppp1r15a^ΔC/ΔC^*lungs, confirming antibody specificity.

### Mice deficient in PPP1R15A induce higher levels of IL-1β and KC in response to cigarette smoke

Acute treatment of mice with acrolein, an oxidising component of cigarette smoke, has been shown to induce expression of PPP1R15A (42). This appeared to contribute to toxicity since *Ppp1r15a^-/-^* mice had better preserved lung architecture when treated with acrolein. To determine the effect of PPP1R15A expression on the proinflammatory signalling elicited by subacute smoke, we exposed wild-type and *Ppp1r15a^ΔC/ΔC^*mice to whole cigarette smoke for 14 days (Figure 7). As we had seen for PPP1R15A protein in ALI-PBECs, prolonged exposure to smoke partially suppressed the expression of PPP1R15A mRNA in mouse lungs (Figure 7A). By contrast, CHOP was unaffected. Smoke increased the protein levels of the pro-inflammatory cytokines IL-1β, IL-17a/f and KC in mouse lungs, and this was significantly higher in the *Ppp1r15a^ΔC/ΔC^*mice (Figure 7). Of note, differences in cytokine levels between the genotypes were not yet manifest after only 3 days of smoke exposure (Supplementary Figure 2). Although smoke also increased the expression of TNFα, GM-CSF, IL-6, IL-10, MCP-1, MIP-1α and MIP-2, these were not significantly different between wild-type and *Ppp1r15a^ΔC/ΔC^*animals (not shown).

**Figure 7.**
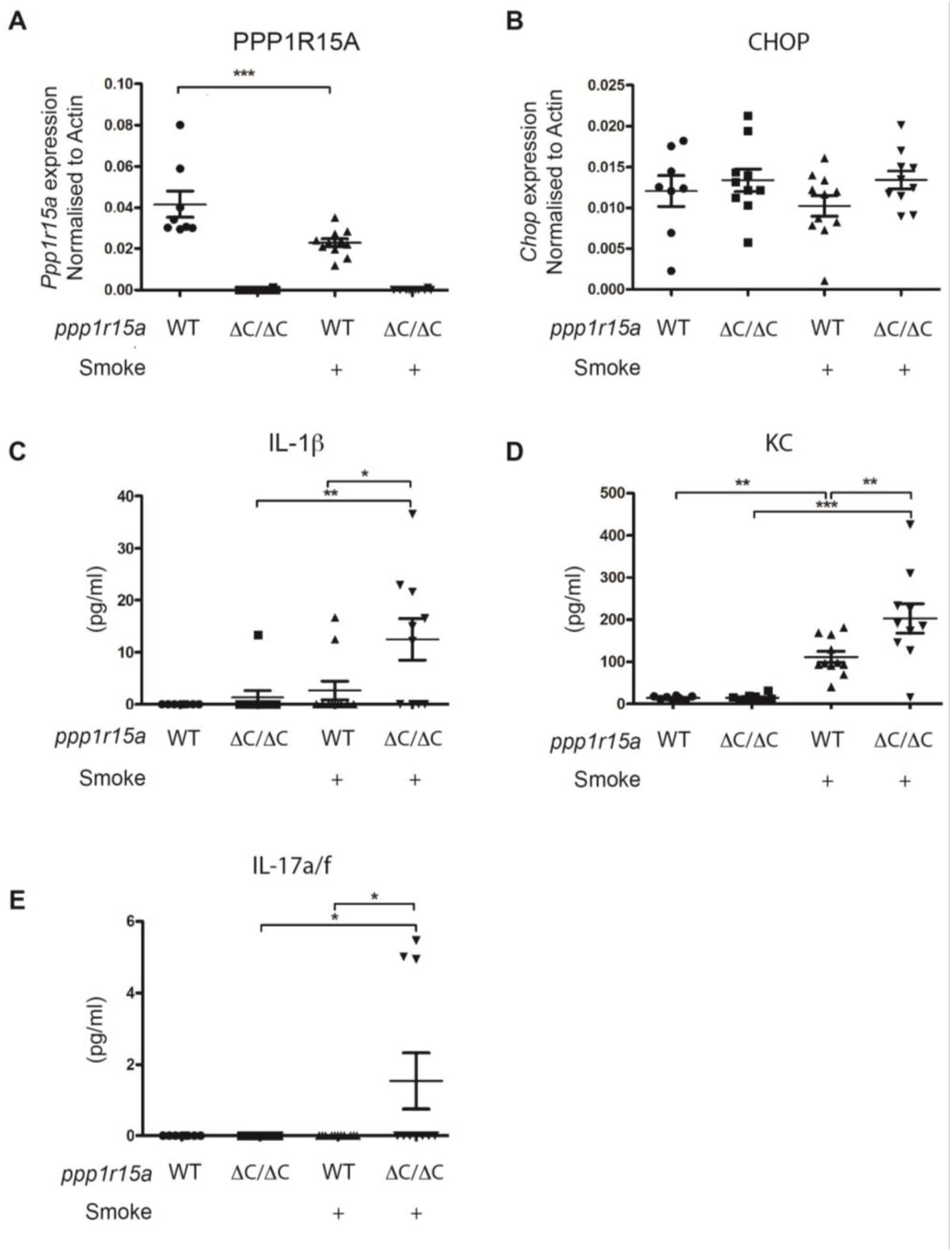
Effects of cigarette smoke exposure on inflammatory cytokine production and stress-response gene expression in WT and *Ppp1r15a^ΔC/ΔC^* mice. (A-B) Expression of *Ppp1r15a* and Chop in whole lung tissue was determined by quantitative RT-PCR and normalised to β-actin. (C-E) Cytokine concentrations in bronchoalveolar lavage fluid (BALF) following 14 days of cigarette smoke exposure. BALF samples were concentrated five-fold using 3 kDa Amicon filters prior to analysis using a Meso Scale Discovery multiplex assay. Cytokines measured included IL–1β, KC and IL–17A/F. Values below the limit of detection were assigned a value of zero. Statistical analysis was performed using one-way ANOVA with Bonferroni’s post hoc test. Data are presented as mean ± SEM. * p < 0.05, ** p < 0.01, *** p < 0.001..

Taken together, our results demonstrate that PPP1R15A is induced by components of cigarette smoke via multiple pathways, including oxidative stress. In vivo, this modifies proinflammatory cytokine secretion.

## Discussion

In this study, we used multiple complementary models to demonstrate that short-term exposure of airway epithelial cells to cigarette smoke activates the ISR. Whole cigarette smoke rapidly caused transient ISR activation in ALI-PBEC. During the recovery period, the kinetics of eIF2α dephosphorylation and recovery of protein synthesis corresponded with induction of PPP1R15A protein expression, suggesting that transient ISR activation by smoke is attenuated by PPP1R15A. However, PPP1R15A induction by smoke was independent of the ISR and was reduced by antioxidant treatment, suggesting the involvement of oxidative stress. In vivo, the absence of PPP1R15A was associated with higher secretion of inflammatory cytokines IL-1β, KC and IL-17a/f in response to subacute cigarette smoke exposure.

We previously showed that acute exposure to cigarette smoke caused transient disruption of the epithelial barrier integrity in ALI-PBECs (18). Furthermore, we showed that cigarette smoke transiently impaired epithelial wound repair by reducing cell migration (25). Interestingly, we found that recovery of epithelial barrier and wound repair functions occurred 6 hours after exposure, which corresponds with the kinetics of PPP1R15A protein expression and the recovery of protein translation it mediates. It is therefore tempting to speculate that PPP1R15A-dependent recovery of protein translation might be responsible, at least in part, for the recovery of ALI-PBEC cultures following smoke exposure. However, we lack conclusive evidence that PPP1R15A expression is cytoprotective in this context.

We observed higher smoke-induced expression of *CHOP* and *PPP1R15A* mRNA in ALI-PBEC cultures from COPD patients when compared to non-COPD controls. Consequently, there was a correlation between expression of these ISR genes and lung function parameters. This is in line with our previous work that revealed an ATF4 target-gene signature in the airway of individuals with COPD, although *PPP1R15A* and *CHOP* were not part of that transcriptome dataset (15). Other ATF4 target genes, such as *PTGS2*, *DUSP5* and *cFOS*, that were found to be increased in COPD airways (15), were not different between COPD and non-COPD cultures exposed to cigarette smoke in this current study. This hints at the existence of additional signalling pathways that act on epithelial cells *in vivo* but do not persist in *ex vivo* cultures. Nevertheless, the more pronounced activation of the ISR in COPD cultures may reflect an increased cell-autonomous susceptibility to cigarette smoke in some individuals or smoke-induced selection of basal cells with an enhanced ISR capacity. This may have inflammatory consequences, as CHOP has been shown to regulate the expression of CXCL8/IL-8 in airway epithelial cells (43, 44).

The chronicity of the stressful insult affects both the nature of the ISR signal and the pathological consequences of the stress (45). This may help explain the differential effects of acute versus chronic exposure to smoke. For example, it is plausible that expression of PPP1R15A may enhance the recovery of protein synthesis that would otherwise be inhibited by cigarette smoke. Indeed, it has been shown that recovery of protein translation mediated by PPP1R15A enables the synthesis of innate immune mediators including IL-6, interferon-α and -β (46, 47), and PPP1R15A also increases GM-CSF translation in a murine model of pulmonary hypertension, exacerbating that phenotype (48). This suggests a pro-inflammatory role for acute induction of PPP1R15A and would account for the reduced lung inflammation observed in *Ppp1r15a* knockout mice treated with acrolein (49). However, in contrast to acute exposure to stress, the repetitive insults that occur in real life may have unanticipated consequences. In this light, it is interesting that after 14 days of smoke exposure, we observed increased expression of proinflammatory mediators in PPP1R15A-deficient mice, which was not apparent following much shorter in vivo exposures. Consequently, inhibition of PPP1R15A might not ultimately have a net anti-inflammatory effect. Recently, loss of PPP1R15A was reported to worsen pulmonary fibrosis following intrapulmonary bleomycin delivery (50). Higher levels of TNFα and a trend towards increased IL-1β were detected in the bleomycin-treated *Ppp1r15a^-/-^* mice. Although the mechanism was unclear, loss of PPP1R15A was shown to render fibroblasts prone to accelerated senescence. Since senescent cells acquire a pro-inflammatory senescence-associated secretory phenotype, including increased secretion of IL-1β and KC (51, 52), it is tempting to speculate that similar mechanisms might contribute to the higher levels of pro-inflammatory cytokines that we detected in the lavage fluid of smoke-exposed mice at late time points. This interpretation is supported by more recent evidence that loss of PPP1R15A can enhance tissue inflammation in other contexts. In a murine model of radiation-induced cardiomyopathy, *Ppp1r15a^ΔC/ΔC^* mice developed more pronounced inflammatory infiltrates and enrichment of inflammatory gene expression programmes than wild-type animals (53). Notably, these effects were independent of CHOP and were not recapitulated by loss of PPP1R15A in bone marrow-derived cells, suggesting that PPP1R15A acts primarily within stressed parenchymal tissues to restrain maladaptive inflammatory responses. These reports, combined with our own, support a broader protective role for PPP1R15A during chronic tissue injury that may explain the augmented cytokine production observed in our smoke-exposed mice. Furthermore, since prolonged cigarette smoke exposure also alters differentiation of cultured airway epithelial cells (54), we cannot formally exclude the possibility that altered epithelial differentiation involving PPP1R15A expression induced by prolonged smoke exposure may in part explain the findings in our mouse study.

An intriguing finding from our study is that PPP1R15A appears to be constitutively expressed in the lung even in the absence of experimental stress. Analysis of our previously published single-cell RNA sequencing dataset (24) revealed that *Ppp1r15a* is the predominant PPP1R15-family transcript expressed in the murine lung, and immunohistochemistry confirmed expression of PPP1R15A protein in airway epithelial cells. These observations suggest that PPP1R15A fulfils physiological functions in the respiratory epithelium beyond simply terminating acute ISR activation. Consistent with this concept, recent work demonstrated constitutive PPP1R15A expression in human bronchial epithelium and identified an antiviral activity of PPP1R15A that restricts influenza A virus replication through its PEST domain independently of phosphatase activity (55). Together with our observation that loss of PPP1R15A exaggerates inflammatory responses to cigarette smoke, these findings support an emerging model in which PPP1R15A acts as a constitutive component of airway homeostasis and host defence rather than solely as an inducible feedback regulator of eIF2α phosphorylation.

Similar to our results with airway epithelial cells, we observed that cigarette smoke condensate increased the expression of CHOP and PPP1R15A in MEFs. This is in line with other observations that suggest that activation of the ISR by cigarette smoke is not restricted to the airway epithelium (8, 56). However, the use of fibroblasts possessing a non-phosphorylatable mutant form of eIF2α, revealed that expression of PPP1R15A in response to smoke can occur independently of the ISR and is abrogated by co-treatment with antioxidants. Similar results were observed with the airway epithelial cell line 16HBE. This is in line with a previous report, indicating that arsenite-induced oxidative stress also increases PPP1R15A expression in an eIF2α-independent manner (57). Previous reports have shown that activation of the p38 MAPK pathway led to increased expression of PPP1R15A (58), and cigarette smoke condensate activates p38 in airway epithelial cells in an oxidative stress-dependent manner (21). More recently, regulation of PPP1R15A has also been shown to occur through post-transcriptional mechanisms. It was observed that cellular stress stabilises *PPP1R15A* mRNA through p38/MAPK/MK2- and PI3K/Akt-dependent phosphorylation of the RNA-binding proteins TTP and Brf1, thereby reducing degradation of PPP1R15A transcripts (59). This provides a plausible mechanistic explanation for our observation that cigarette smoke induces PPP1R15A independently of eIF2α phosphorylation and extends earlier work showing oxidative stress-dependent induction of PPP1R15A via p38 signalling. Thus, cigarette smoke may increase PPP1R15A expression not only through canonical ISR signalling but also through oxidative stress-driven stabilisation of PPP1R15A mRNA.

In summary, we have shown that PPP1R15A is induced in vitro and in vivo by cigarette smoke and modifies inflammatory signalling. This is more pronounced in COPD airway epithelial cells and correlates with disease severity. The absence of PPP1R15A activity during subacute cigarette smoke exposure is associated with higher pulmonary levels of pro-inflammatory mediators.

## Acknowledgements

This study was supported by an unrestricted research grant from Galapagos N.V. and a Fellowship grant (#9.1.14.082FE) from the Lung Foundation Netherlands. SJM was supported by the MRC (G1002610, MR/V028669/1 and MR/R009120/1), EPSRC (EP/R03558X/1), Cambridge Biomedical Research Centre (BRC-1215-20014); British Lung Foundation, Asthma+Lung UK, Royal Papworth Hospital, Alpha1 Foundation.

---

**Supplementary Figure 1.**
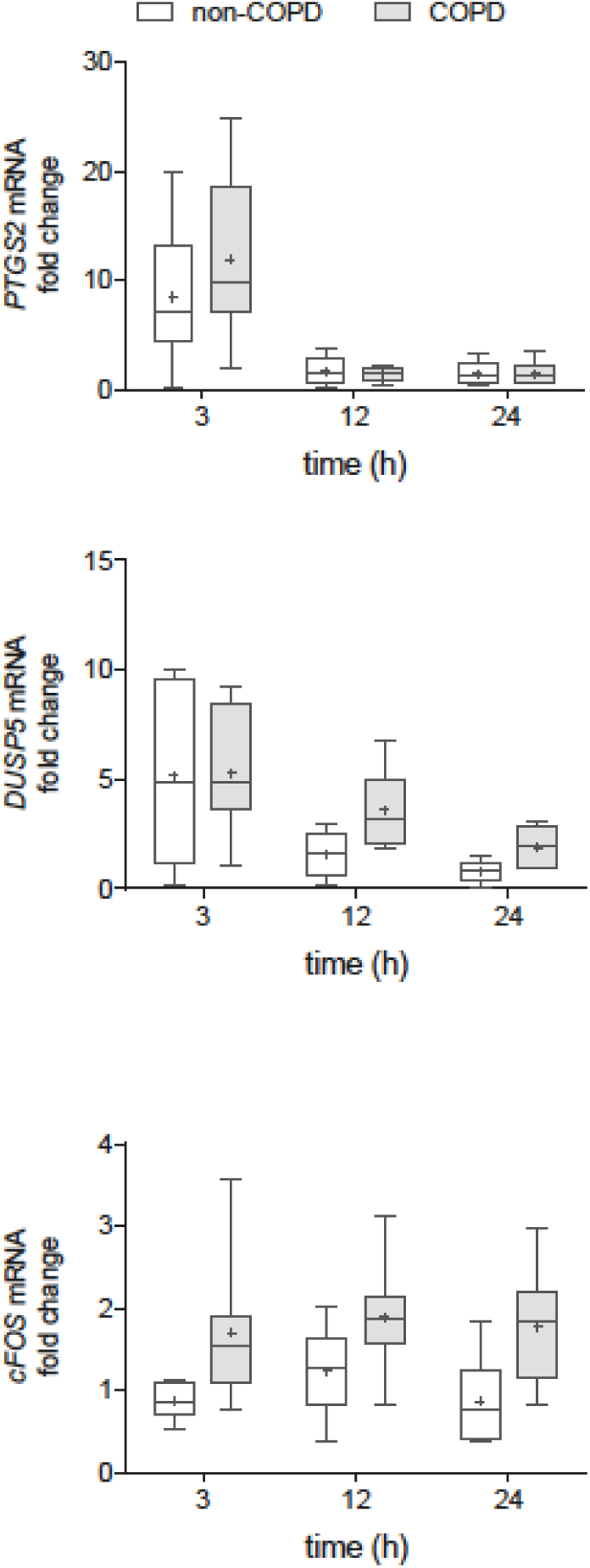
Genes upregulated in COPD lungs by ATF4 that did not differ between COPD and non-COPD ALI-PBECs. CS-induced expression of *PTGS2*, *DUSP5* and *cFOS* in COPD and non-COPD ALI-PBEC cultures. ALI-PBEC from COPD and non-COPD donors were exposed to whole CS and incubated for 3, 12, and 24 h after which *PTGS2*, *DUSP5* and *cFOS* mRNA expression was determined. Cultures from n=7 COPD and n=6 non-COPD donors were used in the analysis. Analysis of differences was conducted using a one-way ANOVA with a Bonferroni *post-hoc* test. Differences between COPD and non-COPD donors were not detected for these genes.

**Supplemental Figure 2.**
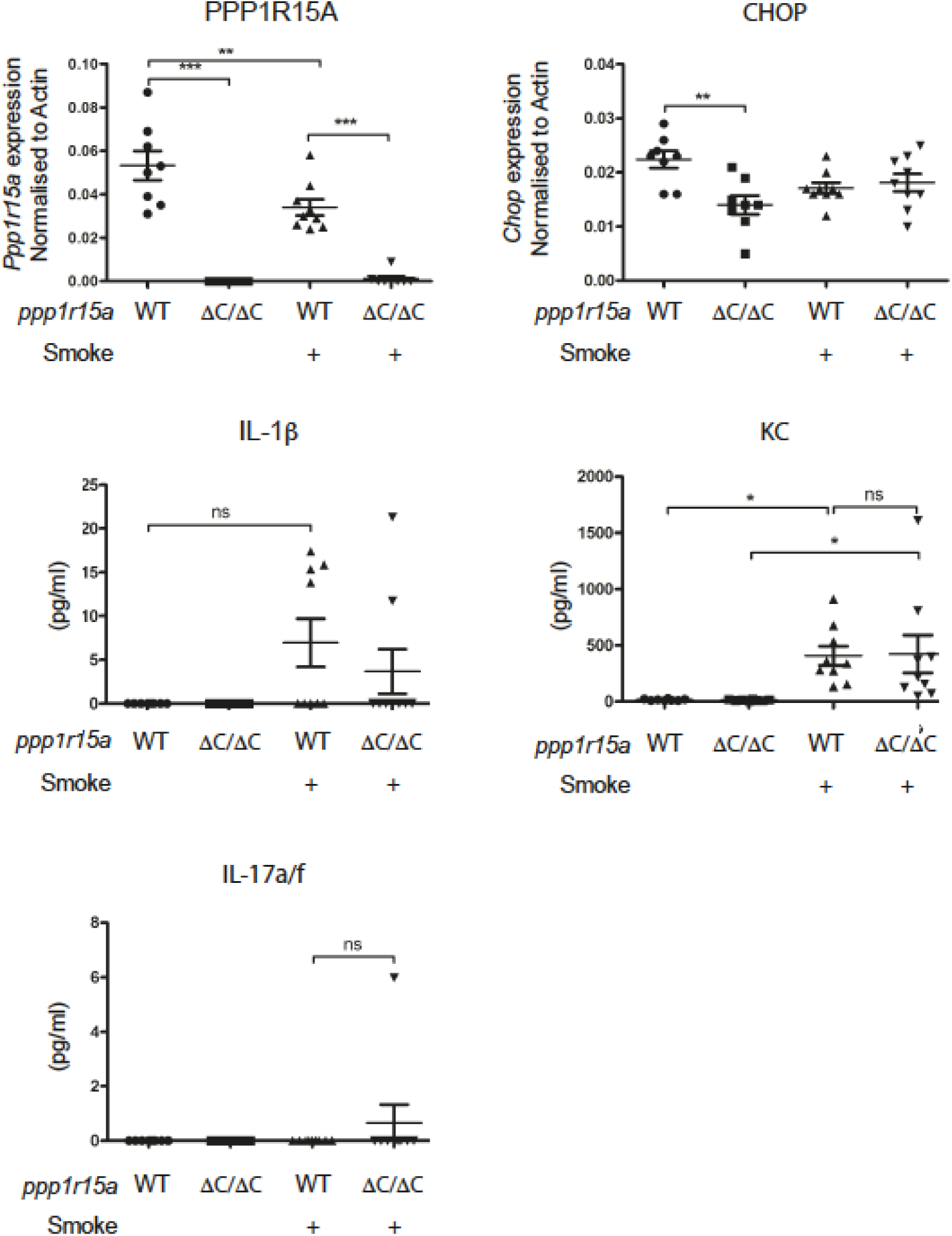
Cytokine levels in bronchoalveolar lavage fluid (BALF) after 3-days of smoke exposure. BALF samples were concentrated (concentration factor 5) follow by Meso Scale Discover ELISA. Results below the level of detection were classified as zero. Analysis of differences was conducted using a one-way ANOVA with a Bonferroni *post-hoc* test. Data are presented as means ± SEM. * p < 0.05, ** p < 0.01, *** p < 0.001.

## Notes

### Competing Interest Statement

The authors have declared no competing interest.

### Summary of Updates

In the revised manuscript we demonstrate the expression of PPP1R15A at the RNA and protein levels in the mouse lung (new Figure 6).

