## Supplementary figures and images for "Cigarette smoke induces PPP1R15A via oxidative stress to modulate inflammatory cytokine production by bronchial epithelial cells"

### Supplementary Figure 1

#### Supplemental figure 1

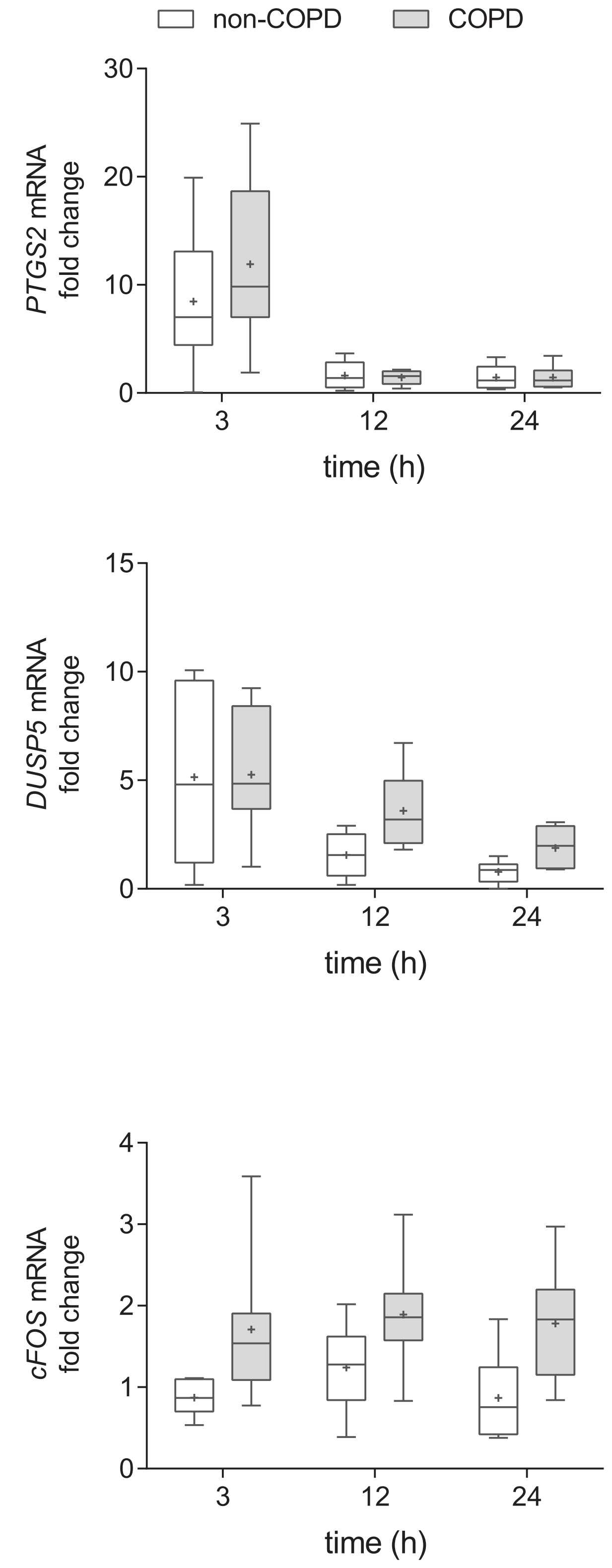

### Supplementary Figure 2

#### Supplemental figure 2

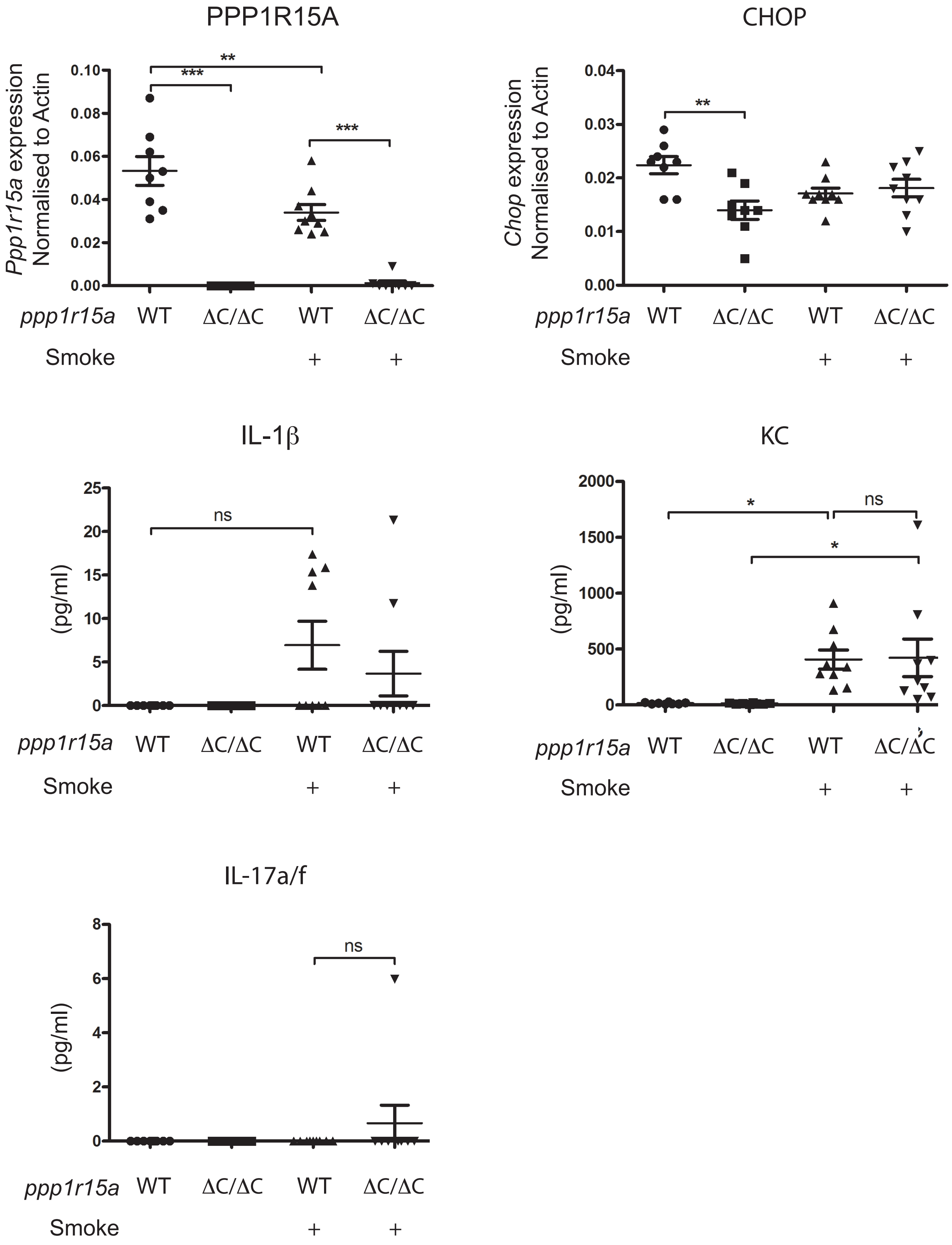
